# Epithelial-Mesenchymal Transition: An Organizing Principle of Mammalian Regeneration

**DOI:** 10.1101/2023.05.11.539303

**Authors:** Kamila Bedelbaeva, Benjamin Cameron, Jack Latella, Azamat Aslanukov, Dmitri Gourevitch, Ramana Davuluri, Ellen Heber-Katz

**Author notes:** **Conflict of Interest:** The authors have declared that no conflict of interest exists.

## Abstract

The MRL mouse strain is one of the few examples of a mammal capable of healing appendage wounds by regeneration, a process that begins with the formation of a blastema, a structure containing de-differentiating mesenchymal cells. HIF-1α expression in the nascent MRL wound site blastema is one of the earliest identified events and is sufficient to initiate the complete regenerative program. However, HIF-1α regulates many cellular processes modulating the expression of hundreds of genes. A later signal event is the absence of a functional G1 checkpoint leading to G2 cell cycle arrest with increased cellular DNA but little cell division seen in the blastema. This lack of mitosis in MRL blastema cells is also a hallmark of regeneration in classical invertebrate and vertebrate regenerators such as the planaria, hydra, and newt. Here, we explore the cellular events occurring between HIF-1α upregulation and its regulation of the genes involved in G2 arrest (EVI-5, γH3, Wnt5a, and ROR2), and identify EMT (Twist, Slug) and chromatin remodeling (EZH-2 and H3K27me3) as key intermediary processes. The locus of these cellular events is highly regionalized within the blastema, occurring in the same cells as determined by double staining using immunohistochemistry and FACS analysis, and appear as EMT and chromatin remodeling followed by G2 arrest determined by kinetic expression studies.

## INTRODUCTION

It is well accepted that mammals do not regenerate appendages whereas amphibians have an impressive regenerative ability (1–3). Besides amphibians, extensive regeneration is also seen in planaria, starfish, sea cucumbers, and hydra (4–6), for example.

A structural key to the regenerative process is the early formation of the blastema which develops after a wound. The amphibian accumulation blastema found in the re-growing limb is a mass of multipotent cells with stem cell markers, thought to be derived from cells migrating into the wound site or from local cells de-differentiating after the process of re-epithelialization and closure of the limb amputation wound. The wound epidermis with the formation of an apical ectodermal cap (AEC) (7) is comprised of active epithelial cells producing factors (8) such as fibronectin and matrix metalloproteinases (MMP) (9, 10). These factors have the potential to affect communication with the mesenchyme, and for this to occur, the basement membrane must be broken down (11). Once that happens, mesenchymal cells begin to accumulate in the space above the cartilage and under the AEC. However, interestingly, little mitosis is seen in the blastema (12).

This lack of mitosis has been a point of interest for the regeneration community for some time. It was shown that blastema cells in the amphibian (axolotls and salamanders) had a high level of DNA synthesis with continuous labeling (approximately 80%) but a very low level of mitosis (around 0.4%) (12–15). A reduction in blastemal mitosis was observed not only in amphibians (12) but also in hydra (16, 17) which have regenerative cells which go through cell cycle entry and then stop in G2/M, and is also seen in planaria (18, 19). In the mammalian liver which is known to regenerate, G2 arrest is seen in adult hepatocytes which are 70% tetraploid (20, 21).

In-vitro studies with newt myotubes showed serum stimulation, cell cycle entry and then G2 arrest (22). This mitotic reduction and G2M arrest has features similar to what we found in the regenerative MRL blastema, and in both normal and blastemal cells in culture but not in non-regenerative C57BL/6 cells (23, 24). Cell cycle analysis revealed that the majority of these cultured cells were found to be in G2M, showed a DNA damage response expressing p53 and γH2AX, and lacked the expression of p21^cip/waf^ protein (CDKN1a), a key G1 cell cycle checkpoint regulator. The lack of p21 in the regenerating MRL mouse predicted that its elimination in otherwise non-regenerating mice would convert these to regenerators. Indeed, p21 KO mouse ear holes could regenerate (close earholes) like the MRL (23, 24).

An obvious question is why or what are these cells doing during G2 arrest without mitosis. Our interest was to identify such cells in the regenerating MRL mouse ear tissue and examine molecules that might reveal information about this. This was the motivation of the current study.

We used markers of G2 arrest (EVI-5, γH3, wnt5a, and ROR2) to identify such cells. We show here that these cells appear beneath the day 7 MRL wound epidermis and in the newly forming blastema but not in B6 tissue. Interesting, these cells were located to a region where EMT seemed to be occurring (a region where the basement membrane breakdown was occurring). Using the EMT markers (Twist and Slug), we found significant overlap between the two populations.

The breakdown of the basement membrane (BM) between the epidermis and dermis, is also a hallmark of amphibian regeneration, and when blocked in the axolotl leads to acute scar formation and a complete cessation of the regenerative response (14, 15). Similar basement membrane breakdown is seen in the MRL mouse ear hole (25), but not in the C57BL/6 scarring response. Major molecules involved in this BM remodeling process include matrix metalloproteinases (MMPs) (10) in both amphibians and the MRL mouse (25). Basement membrane loss has been associated with an EMT response (26).

One possible function of such cells resting in G2/M was DNA repair or chromatin remodeling. We thus used markers for chromatin remodeling (EZH2, H3K27me3) and found that again, there was significant overlap. Since tissue labeling is not always exact, we isolated day 7 blastemal cells from the MRL mouse ear blastema and showed using FACS analysis that over half of the cells showed triple labeling.

Finally, we previously showed that the hypoxia inducible factor or HIF-1a is highly expressed in the day 7 MRL blastema (27, 28). Here, we show that HIF-1a is also a central activator of EMT and chromatin remodeling.

## Materials and Methods

### Animals

MRL/MpJ female mice were obtained from The Jackson Laboratory; C57BL/6 female mice were from Taconic Laboratories. Mice were used at about 8 to 10 weeks of age in all experiments under standard conditions at the LIMR and the Wistar Institute Animal Facilities. Mice were ear-punched and euthanized on days 2, 3, 5 and 7, and ear pinnae removed as indicated and as previously described (27).

### Tissue Preparation, Immunohistochemistry (IHC), and Microscopy

Tissue from hole-punched ears were fixed with Prefer fixative (the active ingredient is glyoxal) (Anatech) overnight, washed in H20, and put in 70% ETOH. Tissue was embedded in paraffin and 5-μm thick sections cut. Before staining, slides were dewaxed in xylene and rehydrated. Antigen retrieval was performed by autoclaving for 20 min in 10 mM Sodium Citrate, pH 6.0. Tissue sections were then treated with 0.1% Triton and nonspecific binding was blocked with 4% BSA (A7906; Sigma) for 1 h. The primary antibodies and matched secondary antibodies used for IHC are shown in **Table 1**.

**Table 1.** Primary and secondary antibodies used for immunohistochemistry (IHC)

|  | 1st antibody |  |  | 2nd antibody |  |  |  |
| --- | --- | --- | --- | --- | --- | --- | --- |
|  | Company | Cat. no. | Dilution | All From Molecular Probe | Company | Cat. no. | Dilution |
| HIF-1α | Abcam | ab2185 | 1:1000 | Alexa Fluor 488 goat anti-rabbit IgG | Molecular Probe | A11008 | 1:200 |
| EVI5 | Millipore | ABN194 | 1:100 | Alexa Fluor 568 goat anti-rabbit IgG | Molecular Probe | A11036 | 1:300 |
| γH3 | Upstate | 06-570 | 1:100 | Alexa Fluor 568 goat anti-rabbit IgG | Molecular Probe | A11036 | 1:300 |
| Wnt5a | R&D | BAF645 | 1:150 | Alexa Fluor 568 goat anti-mouse IgG | Molecular Probe | A11005 | 1:200 |
| ROR2 | Cell Signaling | 4105 | 1:100 | Alexa Fluor 568 goat anti-rabbit IgG | Molecular Probe | A11036 | 1:200 |
| Twist | Santa Cruz | Sc81417 | 1:150 | Alexa Fluor 488 goat anti-mouse IgG | Molecular Probe | A21121 | 1:200 |
| BMI-1 | Santa Cruz | Sc390443 | 1:250 | Alexa Fluor 488 goat anti-mouse IgG | Molecular Probe | A21121 | 1:200 |
| H3K27me3 | Abcam | Mab6002 | 1:100 | Alexa Fluor 488 goat anti-mouse IgG | Molecular Probe | A21121 | 1:200 |
| EZH2 | Thermo-Sc | MA5-15101 | 1:100 | Alexa Fluor 488 goat anti-mouse IgG | Molecular Probe | A21121 | 1:200 |
| EZH2 | Invitrogen | 3210608 | 1:100 | Alexa Fluor 488 goat anti-rabbit IgG | Molecular Probe | A11008 | 1:200 |

For histological stains, tissue sections were treated the same as above and then stained with Hematoxylin (Leica Microsystems, # 3801562) and Eosin (Leica Microsystems, #3801602). For IHC, tissue sections were then treated with 0.1% Triton and nonspecific binding was blocked with 4% BSA (A7906; Sigma) for 1 h. The primary antibodies and matched secondary antibodies used for IHC are shown in **Table 1**, as above. The slides were washed, rehydrated, cleared with Xylene and coverslipped with Permount mounting media (Fisher, SP15-500). Staining was visualized for fluorescent labeling using a fluorescent Olympus (AX70) microscope and a DP74 camera and cellSens software for image analysis, or bright field for H&E staining. This was previously described (28).

For cultured cell staining, primary fibroblast-like cell lines from ear tissue were established from MRL and B6 female mice and then grown in DMEM-10% FBS supplemented with 2 mM L-glutamine, 100 IU/mL penicillin streptomycin and maintained at 37 °C, 5% CO2, and 21% O2. For immunohistochemical staining, fibroblasts were grown on coverslips in DMEM with 10% FBS at 37 °C in a humidified 5% CO2 incubator. The coverslips were rinsed with 1× PBS, the cells were fixed in cold methanol (−20 °C) for 10 min, rinsed with 1× PBS, treated with 0.1% Triton-X100, and then incubated with the appropriate primary and secondary antibodies (**Table 1**). Photomicrographs were produced using the fluorescent microscope (Olympus AX70) and a DP74 camera with cellSens Standard software for image analysis. Confocal images were captured on a Leica TCS SP5 II laser scanning confocal microscope (Leica Microsystems, Inc., Deerfield, IL) using AOBS and sequential scanning with 405, 488 and 561nm laser lines. Individual frames and short z-stacks were acquired at maximum resolution with a 63X 1.4NA objective following Nyquist criteria. Post-processing for maximum projection and noise reduction was done using Leica LAS-AF software and exported to .tif files.

### FACS analysis

Day 7 MRL ear donuts from 2 mm ear punches were generated using a 4 mm punch to retrieve tissue of interest. These donuts were teased apart, treated with dispase and collagenase, and then immediately stained with multiple antibodies without culturing. Cells were fixed in 4% paraformaldehyde for 15 minutes at room temperature and then washed in excess PBS. Permeabilization was achieved by adding ice-cold methanol to pre-chilled cells, while gently vortexing, to a concentration of 90%, then placing cells on ice for 10 minutes. After washing in excess PBS, cells were stained in 100ul of staining buffer (PBS + 2 % FBS) containing PE Mouse SNAI2/Slug (BD Biosciences 564615), EZH2 eFluor 660 (Thermo Fisher 50-9867-82), and Tri-Methyl-Histone H3 (Lys27) PE-Cy7(Cell Signaling 91611S) (**Table 2**) and incubated for 30 minutes at 4°C protected from light. Cells were resuspended in PBS and the fluorescent signals acquired on the BD FACS Canto II. Compensation was performed using UltraComp eBeads (Thermo Fisher 01-2222-41). Cell gates were drawn based on FSC/SSC and doublets were discriminated prior to analysis. Percentages were obtained using FlowJo software.

**Table 2.**
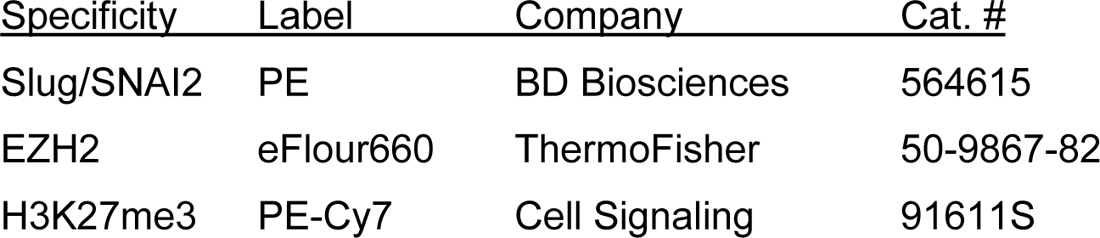
Directly labeled antibodies used for FACS analysis

## RESULTS

### Analysis of Cells in G2M using Multiple Cellular Markers

Through and through 2.1 mm holes were created in the ear pinnae of female MRL and C57BL6 mice. By day 33, the MRL ear hole wounds completely closed with lack of scarring, while the B6 earholes remained for the life of the animal with scarring along the hole perimeter (ref 27, **Fig 1A**). Early in the MRL regenerative response (day 7), the basement membrane has disappeared in the MRL under the wound epidermis but not in the nonregenerative B6 (**Fig 1B, C, arrows**). H&E sections show the distinct border between B6 epidermis and dermis (**Fig 1D, E**) but the lack of cellular organization between the dermis and epidermis in the MRL wound site **(red arrows, Fig 1F, G**) suggesting an ongoing EMT response.

**Figure 1.**
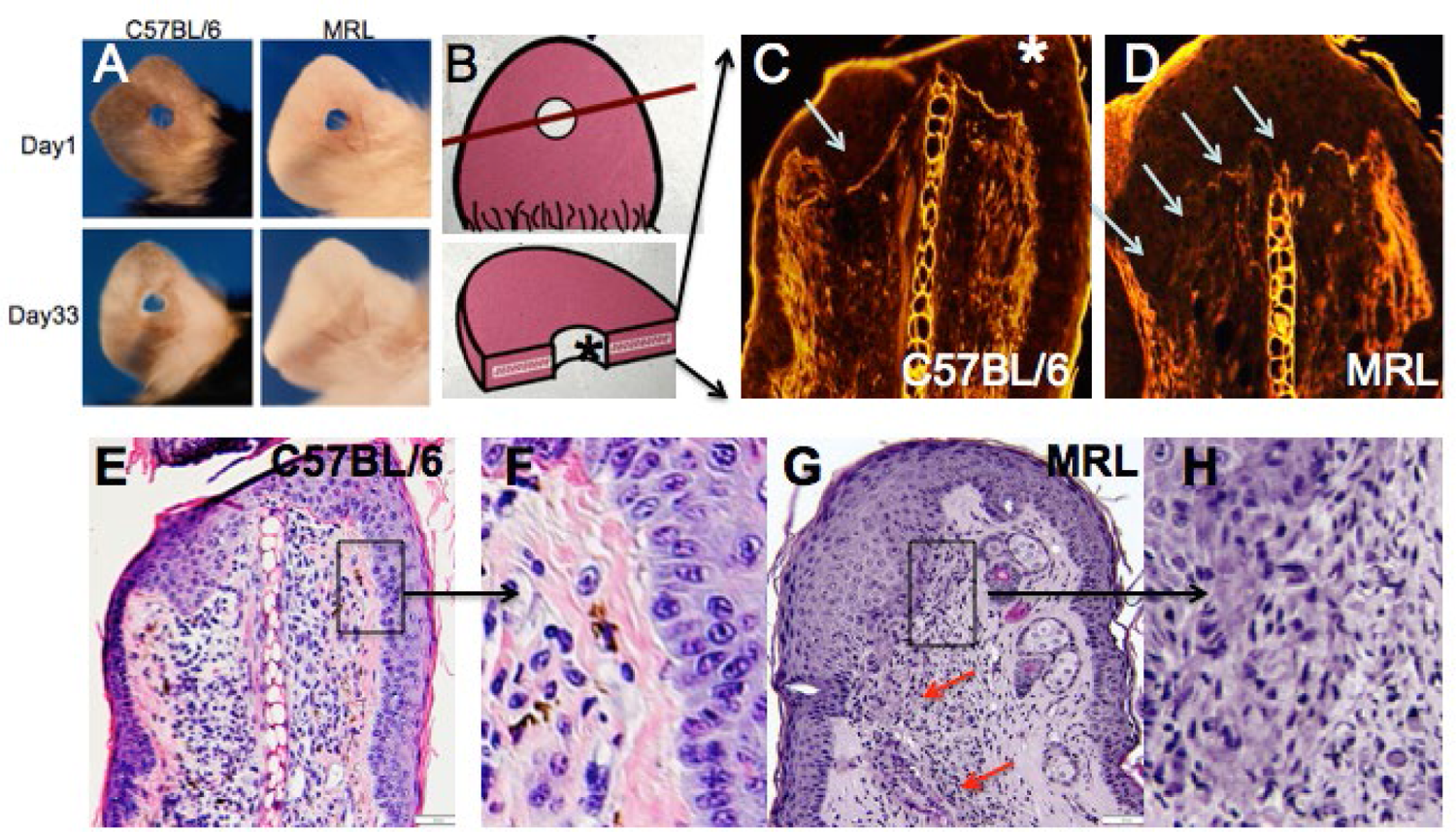
Ear hole closure, basement membrane breakdown, and differences in epithelial-mesenchymal borders on day 7 post ear punching. The MRL mouse when ear punched with a 2.1 mm punch, shows complete ear hole closure (**Fig 1A**) unlike any other mouse strain such as C57BL/6 or B6 (27). The processes have been identified as similar to amphibian regeneration. A diagram of how the ear hole is cut is shown in **Fig 1B**. The upper panel shows the ear pinna with the hole and a line showing how the ear was cut. The panel below shows the ear section with the asterisk indicating the top of the section. One of the early events in both amphibian and MRL regeneration is the breakdown of the MRL but not the B6 basement membrane seen in **Fig 1C, D** (white arrows). Here, injured tissue is stained with H&E and epifluorescence shows stained protein levels and the white line between the epidermis is seen in the B6 but absent in MRL. Examination of the organization of the epidermal/dermal boundary after H&E staining shows a discrete border with a clear and organized basal epidermis in B6, with the boxed area (1E) magnified 3x in (1F), unlike that seen in the MRL boxed in area (1G) and magnified 2x in (1H) with a disorganized and irregular border and no basal epidermis, also seen in other areas of the ear (red arrows). **Fig 1A** is reproduced from (Clark et al. 1998). **Fig 1E** shows a measuring bar = 50 microns and applies to 1C, D, E, and G.

Given the past known deficit in mitotic activity in the blastema along with the finding that cells from the MRL blastema show an unusually high level of G2M arrest (**Fig S1AB**), we questioned where such cells in the healing ear tissue might be found in order to study them further. Consecutive serial sections taken from paraffin-fixed, hole-punched MRL and C57BL/6 ears 7 days after injury were stained for G2M arrest markers including EVI-5 and phospho-H3 (γH3) (**Fig 2A-H**) as well as wnt5a and ROR2, also considered G2M markers (**Fig S1C-F).** These molecules had been previously shown to have increased expression in the MRL ear compared to the B6 ear (24, 28). B6 tissue showed little or no IHC staining for any of the G2M markers, compared to that seen in MRL tissue. MRL and B6 IHC images were overlapped onto H&E images (**Fig 2B, D, F, H)**) from a consecutive slide. MRL IHC for these 4 molecules showed similar but not identical localization, possibly due to differences in the level of the ear wound embedded in the paraffin block. However, a general finding was that there was staining in the area of the epithelial-mesenchymal margin (**Fig S1DF, Fig 2DH).** This staining pattern suggested that we might see co-expression of G2 markers with an EMT marker and this was done by co-staining the same slide for multiple markers.

**Figure 2.**
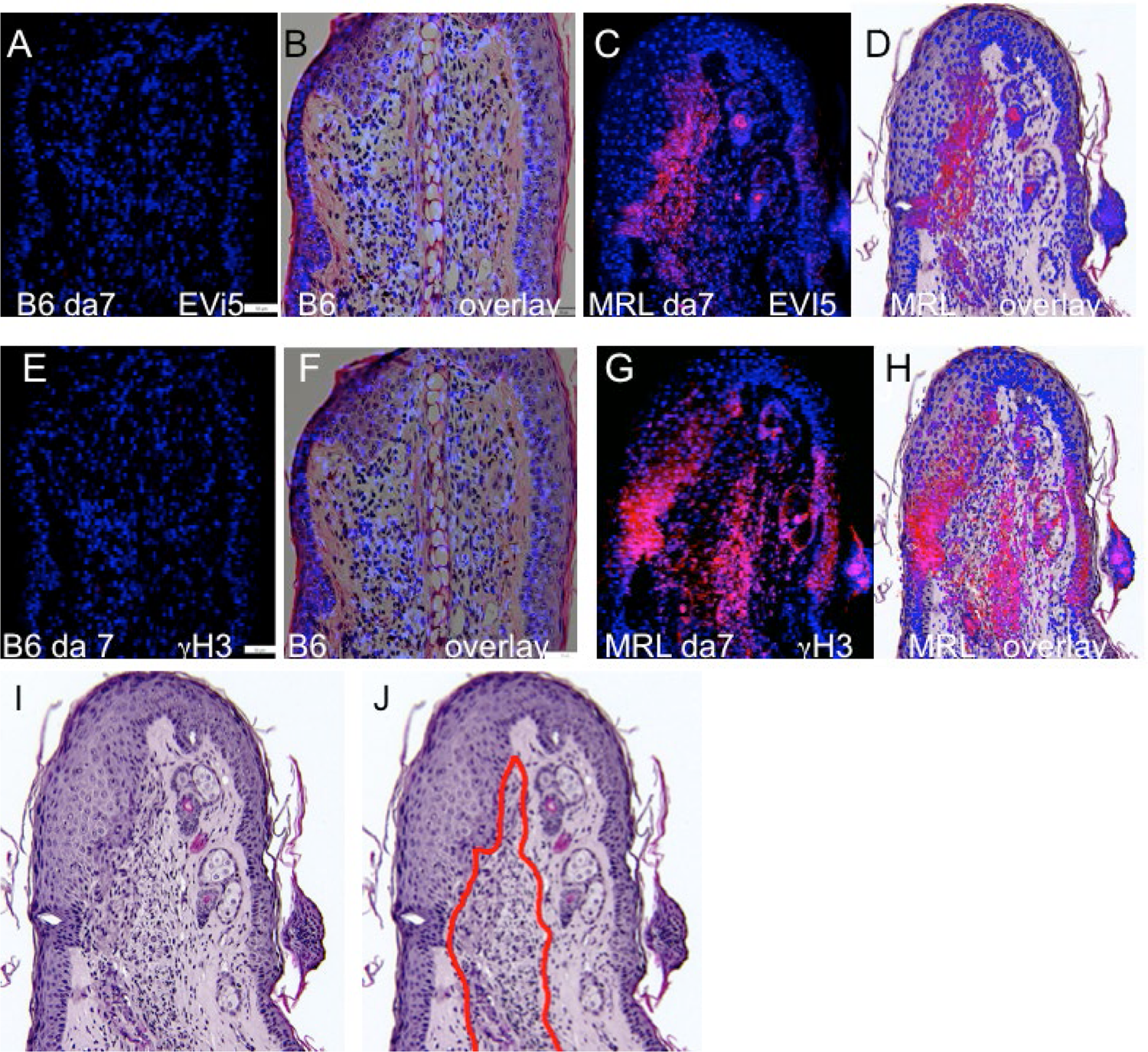
Ear tissue stained with G2M markers. B6 and MRL ear tissues from day 7 post-injury are stained with antibody to EVI5 [**Figs 2A-D**], γH3 [**Figs 2E-H**], and DAPI (**Figs 2A-H**). B6 and MRL stained tissue is overlaid on a consecutive H&E stained section from the same block. Staining for Wnt5a (**Fig S1C, D**), and the Wnt5a receptor ROR2 (**Fig S1E, F**) was also carried out. The zone of reactivity is based on the common staining region as well as cellular changes seen next to the regenerating epidermis in the MRL H&E section [**Fig 2I**] which we have called the “EMT zone” (surrounded by a red line) (**Fig 2J**). Two blocks with ear holes from 2 mice/strain and 3 consecutive sections per slide were stained per antibody. Data from one block for each strain is shown. In Fig 2A and E, the measuring bar = 50 microns but applies to all figures.

### Co-Expression of G2M and EMT Markers in the same Region of the Injured Ear and in individual Cells found there

IHC co-staining of day 7 ear tissue was carried out with both the G2 marker EVI-5 and the EMT marker Twist as seen in **Fig 3**. The locus of Twist (green) staining in the MRL ear day 7 was seen in select regions (**Fig 3bc**). In those Twist-positive regions, EVI-5 (red) showed a similar pattern of staining (**Fig 3ac**). High magnification confocal images show individual cells co-staining for both markers with EVI-5 in and around the nuclear membrane and in the cytoplasm and Twist staining more diffusely in both the nucleus (DAPI+nuclei, blue) and cytoplasm (**Figs 3efg**). These cells were seen throughout the co-stained ear regions.

**Figure 3.**
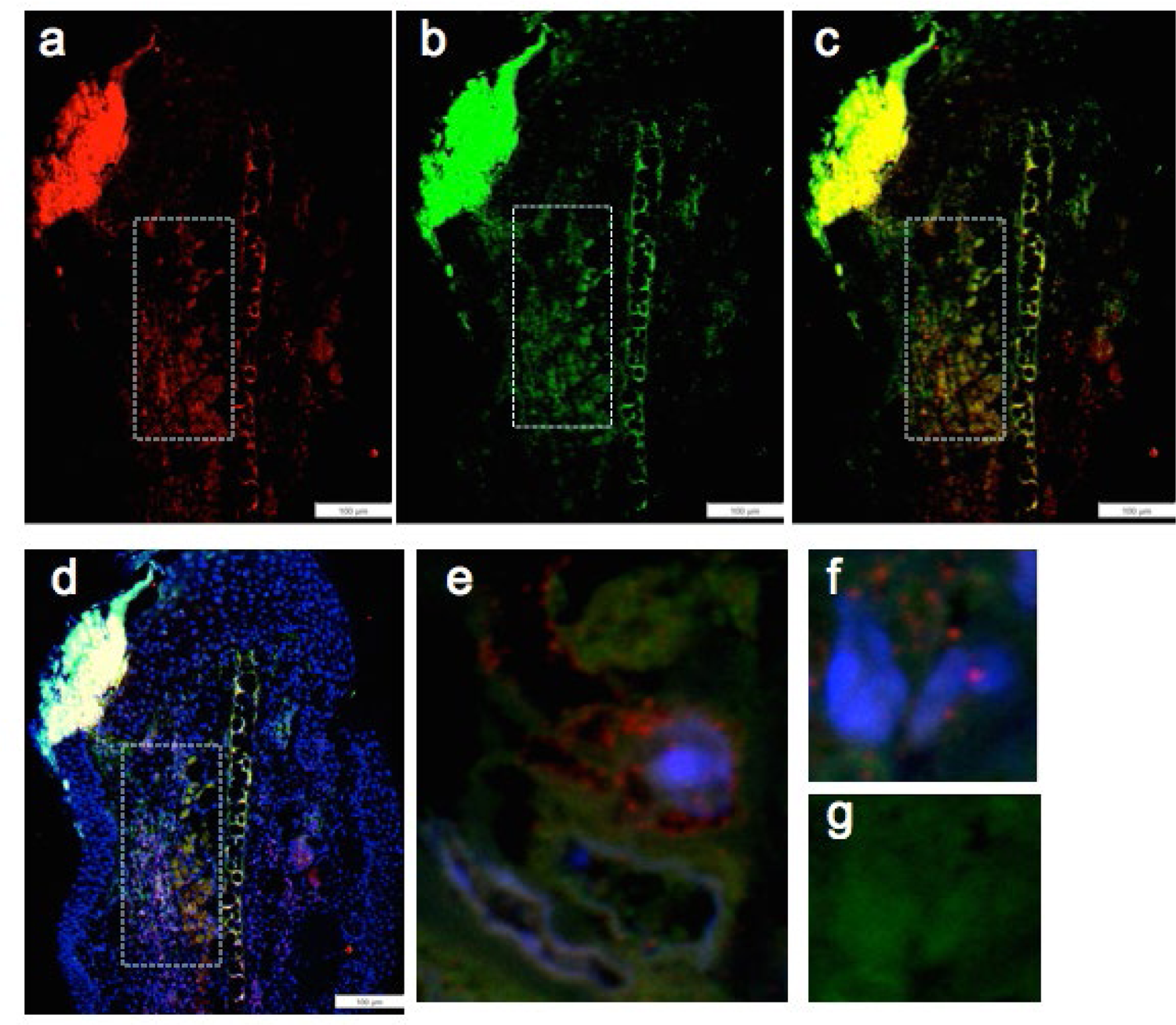
Ear tissue double-stained with the G2M marker EVI-5 and the EMT marker Twist. **Fig 3a-d** shows a comparison of the same region in the MRL ear tissue (white boxes) from day 7-post injury stained with antibody to a) EVI5 (red), b) Twist (green), c) a+b and d) a+b+DAPI (blue) and found in the “EMT zone”. Confocal images are seen in **Fig 3e-g** showing co-staining in both the nucleus and the cytoplasm of individual cells. EVI-5 is known to stain in the nucleus, the nuclear membrane, and the cytoplasm associated with tubulin and the cytoskeleton. Two blocks with ear holes from 2 different MRL mice and 3 consecutive sections per slide were double stained. Data from one block is shown. DAPI staining shows the nuclei. The measuring bar = 100 microns. Note: Regions of intense staining in the upper left corner of Fig 3a-d is due to folded tissue in the slide preparation and is an artifact.

### Co-Expression of EMT and Chromatin Remodeling Markers in Individual Cells

We next asked if the unusually high number of cells paused in G2M in regenerating tissue which co-stained with an EMT marker might also be engaged in chromatin remodeling. We thus examined co-expression of Twist, the EMT marker (green), with a chromatin remodeling marker EZH2 (red) (**Fig 4**). EZH2 shows staining in the same location as Twist (**Fig 4a-d**). Throughout that region, high magnification confocal images show individual cells co-staining for both markers with EZH2 mainly associated with the nuclear membrane and Twist found in the perinuclear, cytoplasmic, and nuclear regions of the cells (**Fig 4e-i**).

**Figure 4.**
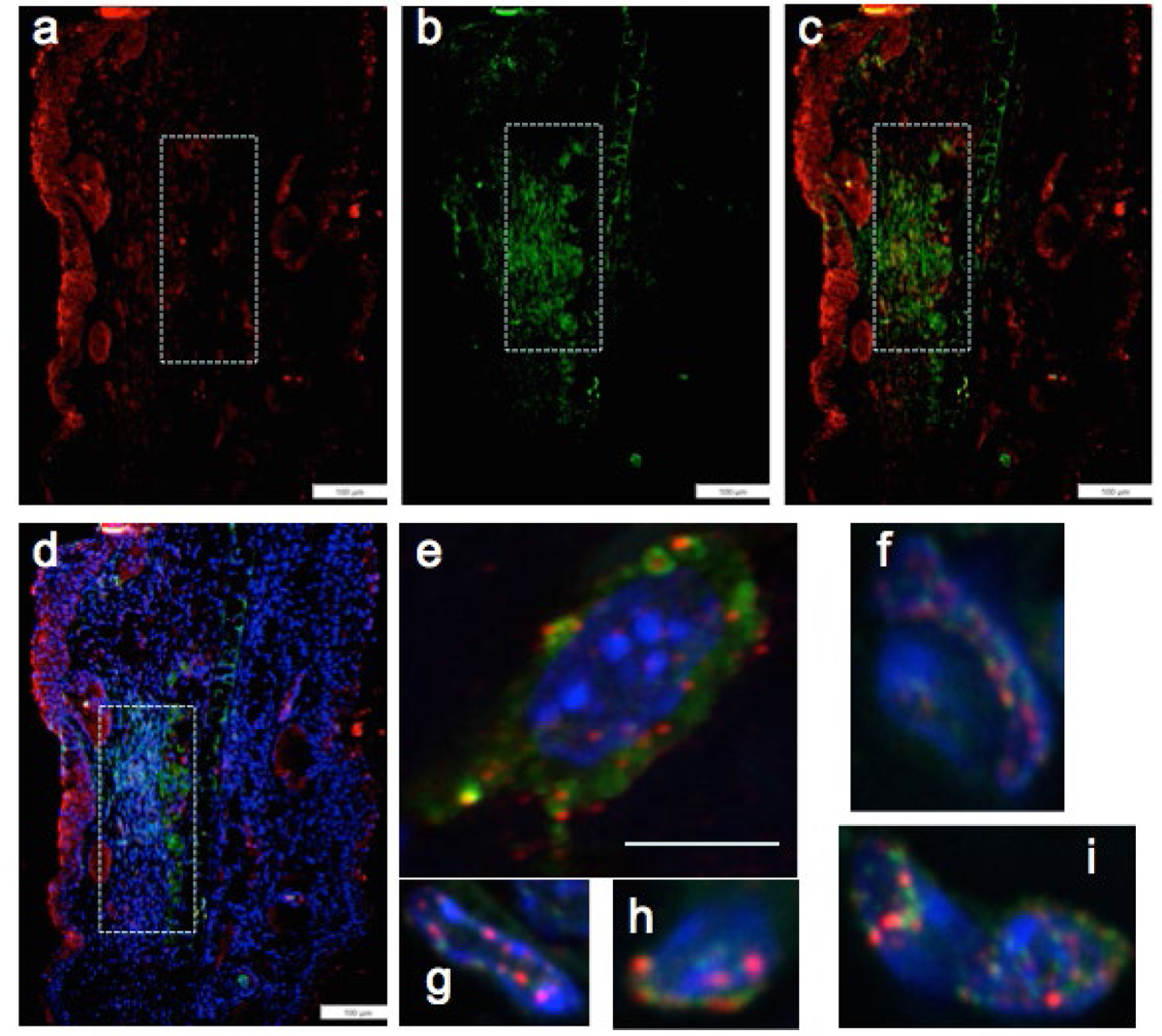
MRL ear tissue 7 days post-injury was analyzed by double staining with an antibody for the EMT marker Twist (green) and for a marker of chromatin remodeling using antibody specific for the PRC2 protein component EZH2 (red). The same area (white boxes) in the MRL blastema co-stained with antibody to a) EZH2 (red) and b) Twist (green), c)a+b, and d)a+b+DAPI again in the “EMT zone”. Confocal images (Fig 4e-i) show EZH2 and Twist antibodies in single cells stained in the nucleus (blue, DAPI), the nuclear membrane, and peri-nuclear region in the same region. Two blocks with ear holes from 2 different MRL mice and 3 consecutive sections per slide were double stained. Data from one block is shown. The measuring bar = 100 microns. The measuring bar for confocal images = 5 microns.

### Co-Expression of G2M and Chromatin Markers in Individual Cells

By pairwise analysis, we tested all possibilities of co-staining. We examined the G2 marker EVI-5 and co-expression with the chromatin remodeling proteins, either EZH2 (**Fig 5A**) or H3K27me3 (**Fig 5B**), the histone H3 which is methylated by EZH2 on lys 27. Staining in both cases was seen in the same area as seen with the previous antibodies (**Figs 5Aa-d, 5Ba-d**). Confocal imaging showed co-staining in single cells throughout the tissue and found in the nucleus and the peri-nuclear region of the cell (**Fig 5Ae, 5Be**).

**Figure 5.**
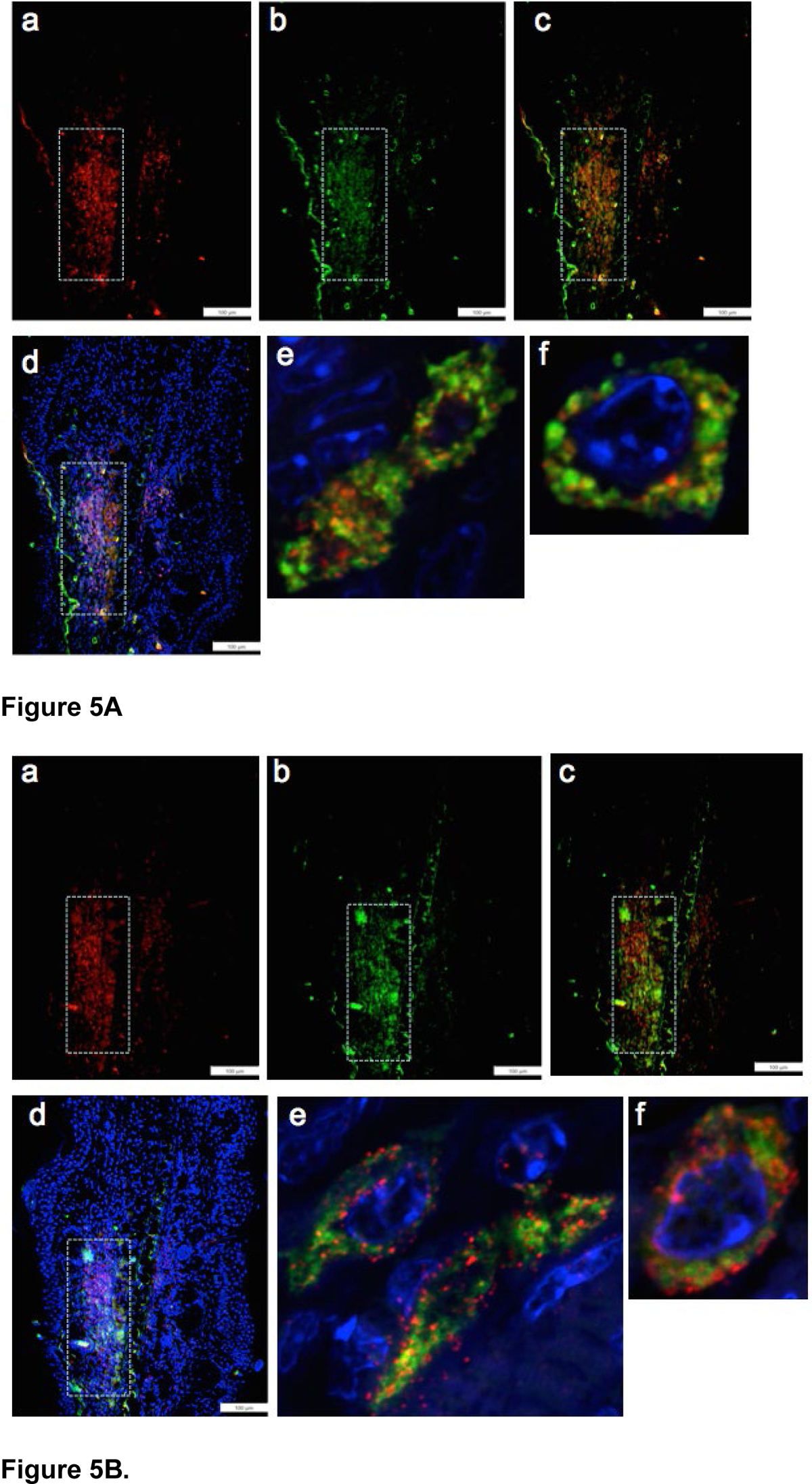
Ear tissue from MRL day 7 post-injury was analyzed pair-wise by double staining with markers for G2M (EVI5) and the PRC2 component EZH2 (**Fig 5A**) and double staining with EVI5 and the chromatin mark H3K27me3 (**Fig 5B**), a product of EZH2 histone transmethylation. Tissue was co-stained with antibodies to a) EVI5 (red), and b) EZH2 (green) (Fig5A) or b) H3K27me3 (green)(**Fig 5B**), c)a+b and d)a+b+ DAPI (blue) to identify nuclei. Areas (white boxes) in the MRL blastema show overlapping staining in the “EMT zone.” Confocal images seen in **Fig 5A**ef and **Fig 5B**ef show same cell staining in the nucleus, nuclear membrane, and perinucleus of EVI5 and either EZH2 or H3K27me. Such single cell staining was seen through the EMT zone. Two blocks with ear holes from 2 different MRL mice and 3 consecutive sections per slide were double stained. Data from one block is shown. DAPI staining shows the nuclei. The measuring bar = 100 microns.

### Pathway Circuit Analysis for all Markers and the Role of HIF-1a

We had previously shown the required upregulation of HIF-1α in regenerating MRL tissue with *siHIF1*α blocking regenerative ear hole closure (28). In **Fig 6A**, we present a gene circuit diagram showing the central role HIF-1α plays in EMT, the chromatin remodeling response, and cell cycle control. Though HIF-1α is extensively expressed throughout the MRL ear blastema on day 7 post-punching injury (and not seen in the B6 ear) (**Fig 6B, C**), the other markers examined here and activated by HIF-1α show a very specifically defined area of expression. This supports the idea that HIF-1α expression is not turning on these genes throughout the whole blastemal region but, in fact, is activating these genes very specifically and regionally (the EMT zone). The numerically labeled pathways are shown in Table 3.

**Figure 6.**
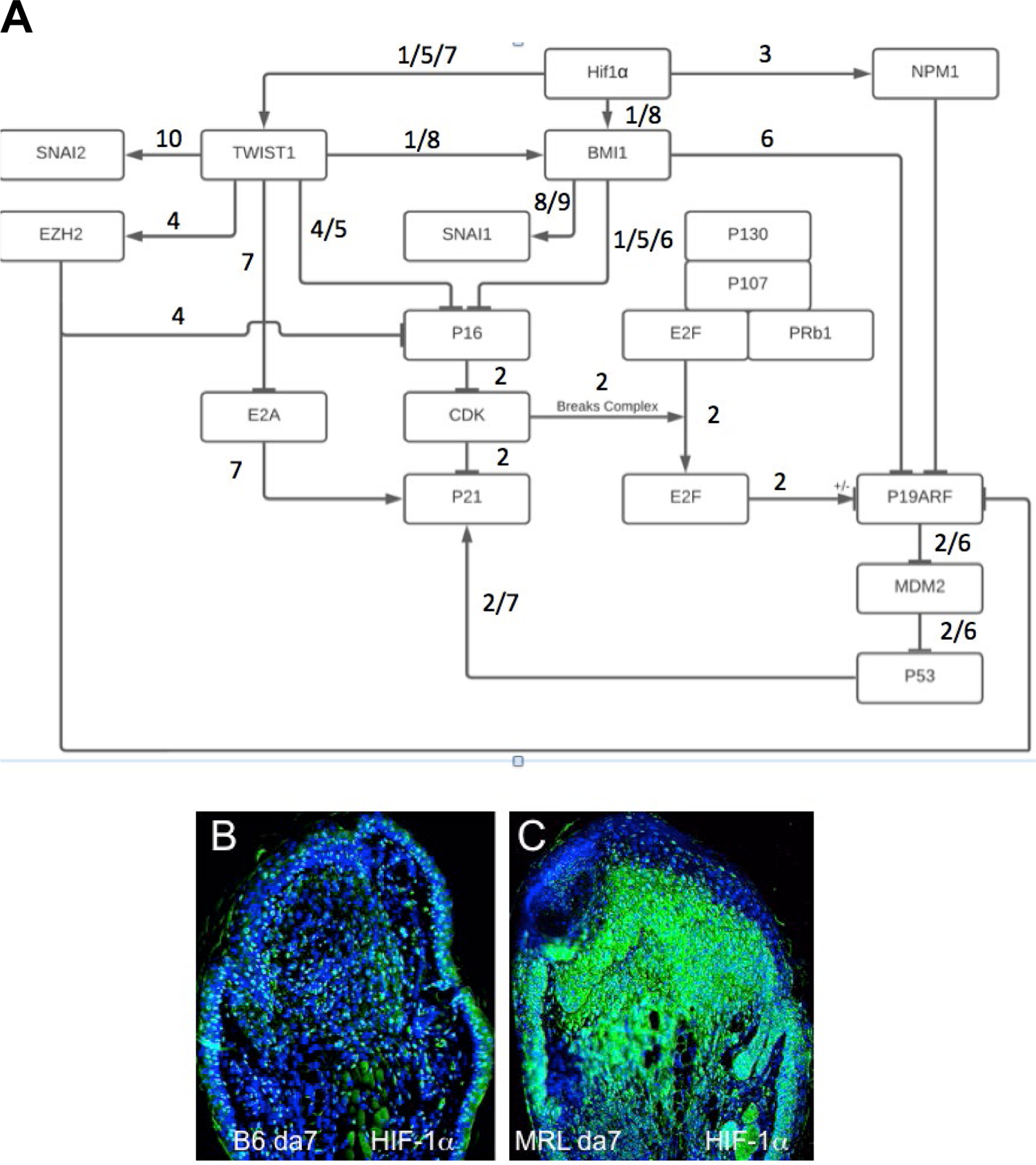
A gene circuit diagram for HIF regulation of EMT, chromatin remodeling, and cell cycle markers. This gene circuit is composed of canonical pathways reference annotated, with HIF-1α as the initiator of down stream events reported herein (**Fig 6A**). HIF-1α has direct activation ties to EMT (Twist) and chromatin remodeling (BMI-1, EZH2 through Twist, and NPM1), and suppression of p19, p21, and p16 cell cycle checkpoint genes. The level of HIF-1α staining in MRL and B6 ear tissue on day 7 shows extensive staining throughout the MRL blastema (**Fig 6B, C**). Four blocks from four mice of each strain with 3 sections per slide were stained with anti-HIF-1α antibody. Only 1 block from B6 and MRL are shown here. The numerically labelled pathways and gene nodes are referenced in Table 3. The measuring bar used in Fig 1= 50 microns applies to these photomicrographs.

**Table 3.** Gene Circuit

### Temporal Expression of HIF-1α/EMT/Chromatin Remodeling and G2M in Injured Ear Tissue

Since the results above were obtained from day 7 post-injury tissue, it was of interest to determine the temporal expression of these markers to understand the sequence of events of these processes. MRL ear tissue was then stained between days 2 through 7 (**Fig 7**). The G2 markers EVI-5 and γH3 were expressed on day 5 but not on day 3 (**Fig 7A-E**), ROR2 was expressed on day 7 but not on day 5 (**Fig 7G, H**). The EMT marker Twist was expressed on day 3 and day 7 but not day 2 (**Fig 7I-K**). The chromatin remodeling PRC1 component BMI-1, and PRC2 component EZH2, and its histone target H3K27me3 were also expressed on day 3 and day 7 but not day 2 (**Fig 7L-Q**). HIF-1α, which directly activates TWIST and BMI-1, was expressed early on day 0 and increased continuously peaking at around days 7-10 by IHC, bioluminescence, and Western analysis (28).

**Figure 7.**
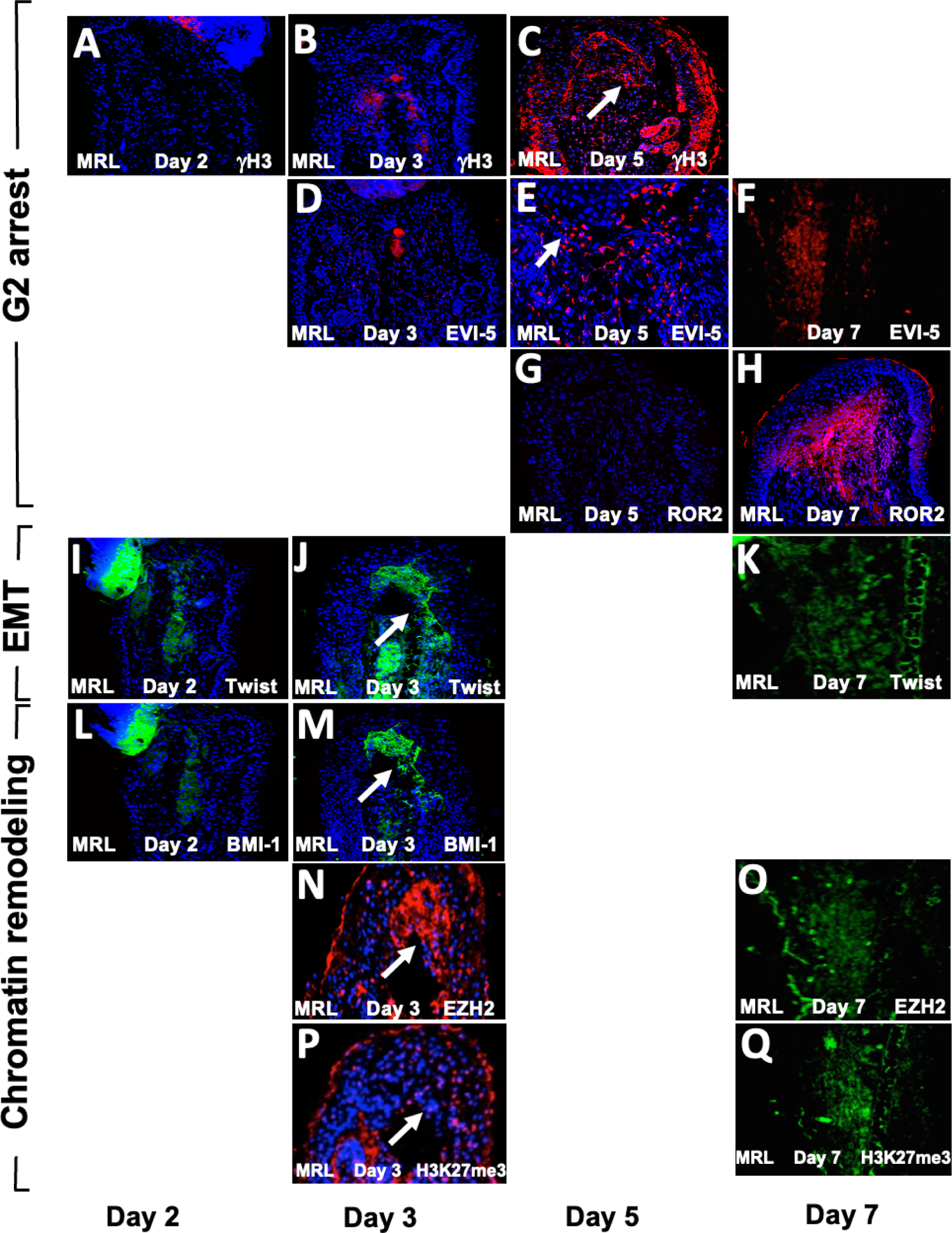
A temporal expression map of G2M, EMT, and chromatin remodeling functional histological markers. MRL ear tissue post-hole punch on days 2, 3, 5 and 7 have been stained with antibody to multiple genes. IHC for G2 genes are seen in **Figs 7A-H**. γH3 and EVI-5 and are expressed on day 5 but not day 3 and ROR2 is not positive on day 5 but positive on day 7. IHC for the EMT gene Twist (**Fig 7I-K**) is positive on days 3 & 7 but not day 2). IHC for chromatin remodeling genes (**Fig 7L-Q**) show BMI-1 which is positive on day 3 but not day 2, and EZH2 and H3K27me3 which are both expressed on days 3 & 7. Areas of interest and IHC positivity are shown by a white arrowhead and more highly magnified micrographs are seen for **Fig 7E** (EVI5 da5) and **Fig 7P** (H3K27me3, da3). The staining in **Fig 7P** looks more diffuse than in other positive figures. The staining seen in some of the micrographs which were not considered positive (**Figs 7A, B, D**) show staining of the cartilage. Two blocks from two injured MRL mouse ears were used for each timepoint with 3 sections per slide. The scale bar in Fig 1, which equals 50 microns applies to **Figs 7A-D, F-Q**. **Fig 7E** is magnified 2x.

### FACS Analysis Shows Markers for all three Functional Sets together in Single Cells

We initially examined MRL and B6 ear-derived fibroblast cells in culture providing higher resolution images of intracellular staining (**Fig 8B**). The majority of MRL cells were in interphase (G2) (**Fig 8Bb**), and did stain for H3K27me3 (**Fig8Bc**), putting G2M together with chromatin remodeling in a single cell and being highly expressed in the nucleus (**Fig 8Bc**). However, B6 cells showed little staining (**Fig 8Ba**).

**FIGURE 8.**
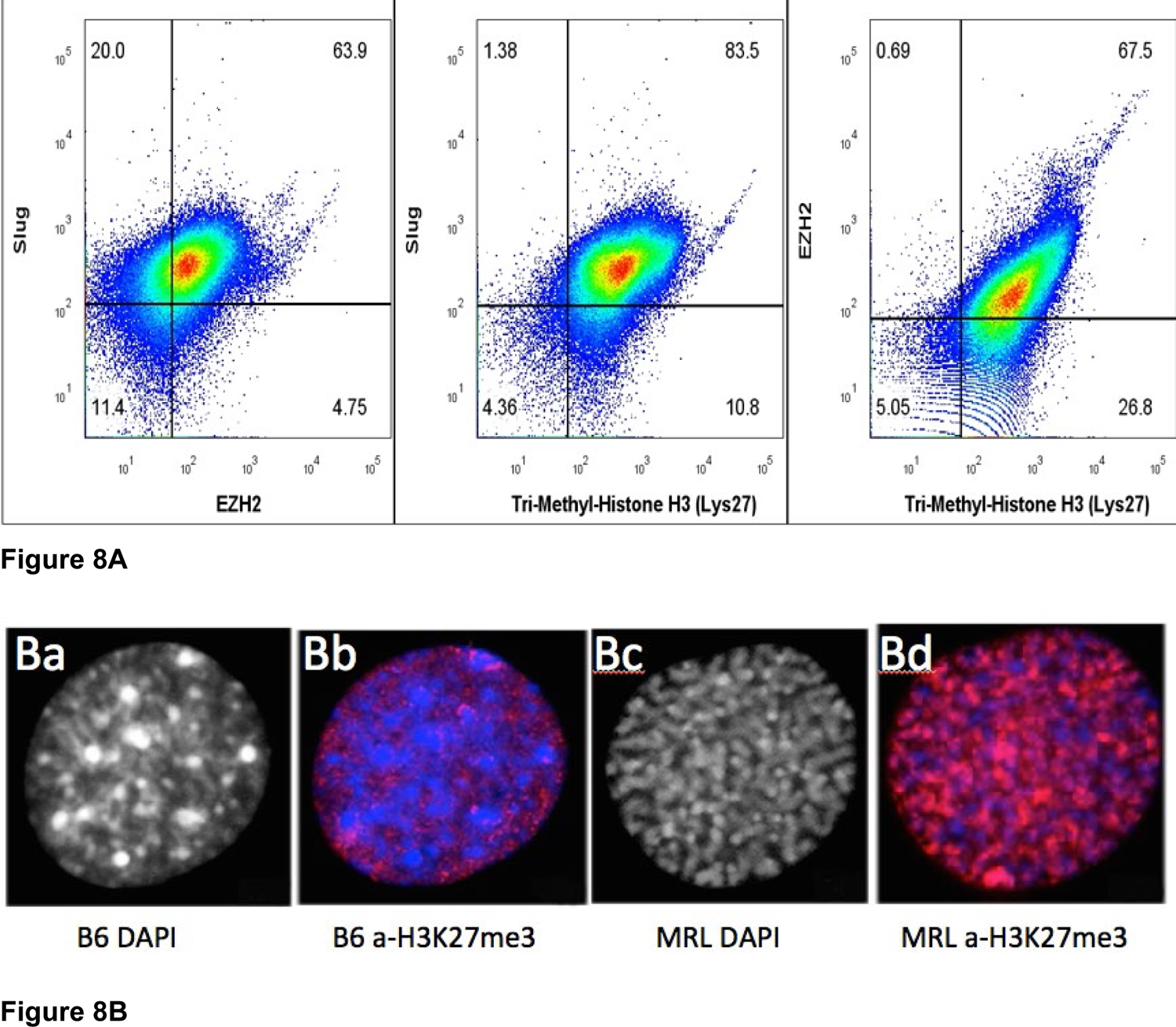
MRL ear tissue-derived cells show the overlap of G2, EMT, and chromatin remodeling markers. Uncultured MRL day 7 cells were stained intracellularly with directly labeled anti-Slug-APC, anti-EZH2-PE, and anti-H3K27me3 PE-Cy7 antibodies (**Fig 8A**). Attempts to label multiple anti-EVI5 antibodies showed no staining. Cells were then analyzed on the BD Canto II and show overlap of the three markers in 79% of the cells (10, 000 events) (**Fig S2**). In a second experiment, to examine the intracellular localization of antibody binding specific for H3K27me3 (red), MRL and B6 fibroblasts were shown be labeled only in the nucleus. B6 showed very faint staining with condensed heterochromatin (prophase, DAPI)) (**Fig 8Ba, 8Bb**) whereas MRL showed strong staining with uncoiled euchromatin (G2, DAPI) (**Fig 8Bc, 8Bd**). DAPI staining in the MRL showed the relationship between G2M and H3K27me3 (**Fig 8Bbc**). Approximately 100 cells for MRL and B6 each were analyzed and showed consistent staining, >85%, for each strain (see **Fig S3**, field of MRL cells).

Since we have not determined the relationship between the cultured cells and the cells present in the ear tissue which we know regenerates, we next used cells derived directly from a day 7 MRL ear-punched pinna. Using 3 different antibodies against Slug (for EMT), EZH2 (chromatin remodeling, and H3K27me3 (chromatin remodeling), a pairwise analysis was carried out using intracellular FACS data of stained cells from ear tissue (**Fig 8A**). There was a significant part of the population, (79%) that co-stained for Slug (EMT), EZH2 and H3K27me3 (chromatin remodeling). Unfortunately, an antibody for G2M (EVI-5 or γH3) to be used for FACS analysis could not be identified. However, with DAPI staining, over 85% of cultured MRL cell nuclei were shown to be in G2 (i.e.uncoiled chromosomes) as opposed to cultured B6 cells which showed condensed chromosome (prophase) (**Fig 8B, Fig S3**).

## DISCUSSION

In our studies of mammalian regeneration in the MRL/lpr and MRL/MpJ mouse, complete scarless closure of a punched hole in the ear pinna occurs within 30 days with cellular and tissue events mirroring that seen in amphibian limb regeneration (27). This includes rapid re-epithelialization, seen in the amphibian within the first 12 hours, and seen in the MRL mouse within 24-48 hours compared to the non-regenerator C57BL/6 mouse which takes from 5-10 days (27). A second hallmark of amphibian regeneration, also seen early in the MRL regenerative response, is the breakdown of the basement membrane (BM) between the epidermis and dermis permitting the cellular and molecular exchange of factors.

The reduction in blastemal mitosis observed not only in amphibians (12, 13) but also in hydra (16, 17) and planaria (18, 19) is a third hallmark of regeneration. This mitotic reduction and G2M arrest has features similar to what we found in the MRL blastema, and in both normal and blastemal cells in culture (23, 24). Cell cycle analysis revealed that the majority of these cultured cells were found to be in G2M, showed a DNA damage response expressing p53 and γH2AX, and lacked the expression of p21^cip/waf^ protein (CDKN1a), a key G1 cell cycle checkpoint regulator. The lack of p21 in the regenerating MRL mouse predicted that its elimination in otherwise non-regenerating mice would convert these to regenerators. Indeed, p21 KO mouse ear holes could regenerate (close earholes) like the MRL (23, 24).

Until very recently (29, 30), EMT, a core process in developmental biology, has received a paucity of attention in the context of regeneration with a notable exception (31). The overlapping localization of cells involved in the processes described above supports the notion that EMT may be the source of functional cells in the blastema participating in the process of de-differentiation, with in-migrating cells acting as bystanders or a supportive milieu. The observation of EMT concurrent with regeneration in organisms spanning evolution from echinoderms to mice suggests a deeper role for this developmental process.

While the studies presented herein are strictly confined to the MRL and C57BL/6 strains of mice which represent what occurs during regeneration (MRL) vs what occurs during wound repair (C57BL/6), we have made repeated comparisons to classical regenerating species including newts and axolotls, which are superior limb regenerators, and to the echinoderm species such as sea cucumber which displays the ability to completely regenerate its gut. The past and ongoing studies in these species are a constant source of insight into mammalian studies.

### Regenerative Epidermis

During the regenerative process, the epidermis plays a very special initiating role. It receives injury signals; it covers the wound and does so rapidly. In the amphibian it occurs within 12 hours and in the MRL mouse regenerating ear hole it occurs within 24-48 hours (27). It forms an apical epithelial cap (AEC), the epithelial structure that covers the wound but has a basal layer which expresses the mesenchymal marker fibronectin (FN) (32, 33). In the normal MRL mouse epidermis pre-wounding, Keratin 16, a gene associated with a keratinocyte activation state, is present at high levels and is not seen in the non-regenerating B6 epidermis (34).

### EMT

Epithelial to mesenchymal transition is a process that is key to developmental events taking place in the embryo for neural crest formation, myogenesis including the heart, gastrulation, stem cell trait acquisition and function, and also tumorogenesis and metastases (35–39). It has been reported to be important in regenerative processes in the sea cucumber and the axolotl (29–31). Though EMT is seen during wound repair, it a very small and transient response within the first 24-72 hours post injury at least in the B6 mouse ear. In the regenerative process, however, as seen here in the MRL mouse ear, it is a full-blown response at least up to day 7. There are multiple stimulators of EMT including hypoxia and molecular activation through molecules such as TGFβ, wnts and others (38).

In this process, epithelial cells, which normally express e-cadherin, cytokeratins, laminin, syndecan, claudin, and desmoplakin, are not motile. After breakdown of the basement membrane in local tumor tissue, for example, they undergo changes by losing their polar epithelial characteristics, such as the expression of e-cadherin, and gain a migratory phenotype, acquiring fibroblast markers such as vimentin, fibronectin, FSP-1, Snail, Slug, Twist, αSMA-1, FOXC2, ZEB1 and N-cadherin (38).

Changes in the basement membrane (BM) are due to MMP activation and remodeling; and this is regulated by HIF-1α. In the amphibian, after limb amputation, the BM does not reform between the epidermis and dermis due to continuous breakdown during the regenerative response. In the mammalian regenerative MRL mouse, MMPs are also activated by HIF-1α, and enzymatically active MMPs are seen as early as day 1 (28). BM breakdown can be viewed as the first permissive step for EMT because it is hard to imagine how any direct cell-cell contact or soluble factor diffusion can occur in the presence of an intact BM.

The MRL BM, after injury, appears to reform on day 4 post-injury, remains intact until day 5, and then again disappears. It may be that on day 4, the BM still allows cell crosstalk in the MRL due to microbreaks in the BM as reported elsewhere (40). Thus, BM breakdown may be happening before, during, and after EMT begins (EMT markers are upregulated on day 3) (**Figs 2, 6**) (41, 42). In the case of the B6 mouse strain, neither MMP expression nor BM breakdown is seen.

EMT is dependent on HIF-1α upstream (43, 44), but it is also dependent on the molecule Twist, a direct HIF-1α target (40, 44, 45), which turns off e-cadherin in mature epithelial cells. Interestingly, the molecule BMI-1 (a direct target of Twist), is found in stem cells, resulting in the de-differentiation step so well known in the formation of the blastema. BMI-1 is a component of the chromatin remodeling PRC1 complex and thus is permissive for the process of chromatin remodeling to proceed. Knockdown of BMI-1 prevented changes induced by Twist and HIF-1α, leading to a lack of expression of stem cell markers (46, 47).

### Chromatin Remodeling

Besides EMT driving chromatin remodeling, the reverse is also true (46–49). Four very different markers of chromatin remodeling are upregulated in the MRL blastema in the EMT zone: (i) The polycomb repressive complex 2 (PRC2) contains the transmethylase EZH2 and is involved in silencing gene expression by (ii) methylating histone H3 on its lysine 27 (H3K27me3) as shown by a specific antibody and (iii) PRC1 binds to and blocks nucleosomes and limits transcription factor access using H3K27 to inhibit RNA pol2 initiation. This complex includes BMI-1, which we show is upregulated in the MRL ear on day 3. Lastly, (iv) nucleophosmin (NMP), a histone chaperone, is found in the nucleolus and binds to H2A and H2B (data not shown). Thus, EMT directly activates chromatin remodeling and there are positive and negative feedback loops.

Though not included in the data presented here is another molecule HDAC3 involved in chromatin remodeling. This is a histone deactylase which has enhancer activity affecting gene expression, is involved in modulating chromatin structure in the nucleus, and is expressed on day 3 post-injury in the MRL. Previous mapping of ear hole closure-associated genes showed that HDAC3 is a strong candidate and is up-regulated in the MRL mouse (50).

### Temporal Sequence

The day 7 IHC results of pairwise staining analysis as well as the days 2, 3 and 5 data shows that markers of all three processes are present throughout the timeframe of early blastema formation. These IHC studies also suggest the temporal order of EMT and chromatin remodeling as being the earliest events occurring with G2 following. However, all three processes co-exist pairwise within the same cells suggesting that all of the expression curves overlap to some degree.

### In-Vitro Studies

From histology, we observed that not all cells in any given region co-stain and there are some regions of the ear blastema where no staining is seen. For in-vitro studies, we isolated fibroblasts from normal unwounded ears from B6 and MRL mice cultured without further selection. The MRL cells showed a predominance of cells in G2M, unlike the B6 cells which were predominantly G0/G1 as shown previously (20). Here, FACS analysis of MRL fibroblasts showed co-straining with SLUG, an EMT marker, and EZH2, a chromatin remodeling marker. However, better resolution could be seen with direct IHC of cultured cells and the intracellular localization of molecular markers is readily apparent. Using an antibody to H3K27me3, we saw co-staining for DAPI showing that the cells were in G2 and interphase, with the same cells expressing high levels of modified chromatin.

One might ask why these normal MRL cultured fibroblasts express markers not seen until after injury in the tissue. Given that the cells are isolated from tissue, are no longer effected by contact inhibition, removed from a normal in-vivo cellular milieu, grown on plastic and selected for growth over time, this might account for early expression of markers.

### HIF-1α and Metabolism

An early marker of regeneration is the upregulation of HIF-1α expression. It was first identified because of the metabolic state used by the MRL mouse which is more embryonic (51, 52) employing aerobic glycolysis with increased lactate in preference to OXPHOS, a shift known to be regulated by HIF-1α. Another clue to the role of HIF-1α was the identification of RNF-7 (50, 53), part of the HIF-1α degradation pathway from MRL gene mapping studies. Also, the drug dichloroacetic acid (DCA), a small molecule inhibitor of PDK1, allows pyruvate to enter the mitochondria to produce ATP shifting metabolism away from glycolysis and towards OXPHOS (52). This blockage of aerobic glycolysis by DCA also blocks regenerative healing in the MRL mouse. Confirmatory evidence for the critical role of HIF-1α was shown by: (a) Blockage of HIF-1α using *siHif1*α leads to blockage of regeneration, and (b) Up-regulation of HIF-1α in otherwise non-regenerating mouse strains converts them to regenerators indistinguishable from the MRL (28). The latter was achieved using of the PhD inhibitor, 1-4-DPCA, in a timed-release hydrogel formulation which down-regulates the degradation of HIF-1α. In addition to ear hole closure, other regenerative models such as enhanced and more rapid liver regeneration (54), and the complete recovery of lost alveolar jawbone (55, 56) as a consequence of periodontal disease are treatable with 1, 4-DPCA. *SiHIF1*α completely blocked ear hole closure showing a HIF-1α requirement in this response (28). We show that HIF-1α is actually the central modulator of the genes examined here and is associated with EMT, G2M arrest, and chromatin remodeling.

### HIF-1α and Cell Cycle Regulation

In the MRL regenerative response, we see that HIF-1α also directly activates Twist and EMT, BMI-1, EZH2 through Twist, NPM-1, and chromatin remodeling. HIF-1α also indirectly suppresses p19, p21, p16 and cell cycle checkpoints. This answers our original question as to why there is so little mitosis in the blastema, and it also ties together important events during regeneration, EMT, chromatin remodeling, and cell cycle regulation. But, why is G2 arrest so critical to regeneration as seen by the fact that the p21 KO mouse recreates the MRL regenerative phenotype? A possible answer is that the p21KO, in fact, upregulates HIF-1α. One interesting possibility is that lincRNA-p21, inhibits HIF-1α and when lincRNA-p21 is off, HIF-1α is up (57). This requires further exploration. From our IHC results, we show that EMT and chromatin remodeling are occurring at approximately the same time, followed by G2 arrest. There are of course developmental studies suggesting this order such as EMT (58) leading to chromatin remodeling which leads to cell cycle changes (59, 60). Reversing chromatin remodeling through the phosphorylation of EZH2 with the resultant inhibition of H3K27 methylation by the cell cycle checkpoint kinase CDK1, which acts in G2 and induces mitosis, leads to differentiation of stem-like cells, providing a possible mechanism for ending the blastema phase of regeneration (61).

## Conclusion

The striking similarity of biological processes that occur at the site of a wound in regeneration-competent species across phyla suggest an evolutionarily conserved organizing principle. A close dissection of the MRL mouse blastema using established molecular markers and their temporal order of appearance identify the Epithelial-Mesenchymal Transition process following HIF-1α expression as a strong candidate for this organizing principle.

## Abbreviations

BMI-1: B cell-specific Moloney murine leukemia virus integration site 1

PRC1: component

EMT: epithelial to mesenchymal transition

EVI-5: ecotropic viral integration site 5 protein homolog

EZH2: enhancer of zeste homolog 2

IHC: immunohistochemistry

PRC2: component

H3K27me3: methylated histone 3 lysine 27

HIF-1α: hypoxia-inducible factor 1α

PRC: polycomb repressive complex.

## Acknowledgements

We would like to dedicate this paper to the memory of Dr. David L. Stocum, retired Emeritus Professor at IUPUI and recently deceased. Dr. Stocum was a giant in the field of regeneration biology and a mentor to many who continue in his footsteps. This work was supported by the grant DE021104 from the NIDCR (NIH), the grant PR180789: W81XWH-19-1-0467 from DOD and partially by the Green Family Foundation. All confocal imaging was carried out by James Hayden. RBP, FBCA, Managing Director/Imaging, The Wistar Institute.

## Author Contributions

EHK designed the experiments, interpreted the data, and prepared the manuscript. KB carried out all immunohistochemistry and microscopy studies, DVG carried out ear-punching and provided all tissue both unstained and H&E-stained slides at the Wistar Institute and AA carried out similar experiments at LIMR. JL generated the pathway circuits. BC carried out the FACS studies. RD provided chromatin remodeling expertise. All authors have read and approved the manuscript.

## SUPPLEMENTAL FIGURES

**Figure S1.**
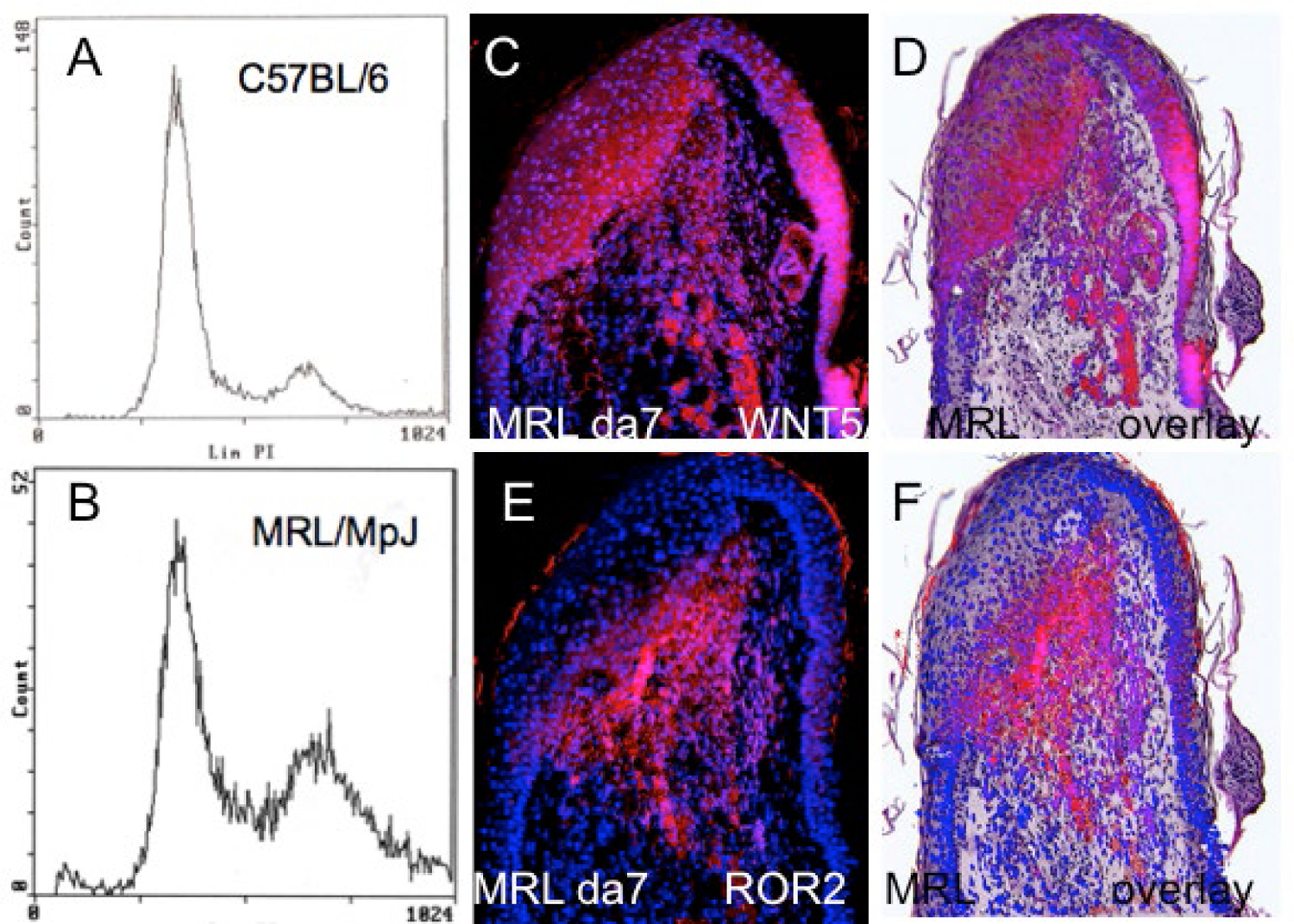
Fibroblasts from MRL ears showed high accumulation in G2M. Cells from C57BL/6 ear pinnae (**S1A**) and from MRL ear pinnae (**S1B**) were examined using propidium iodide staining and FACS analysis to determine cells in different stages of the cell cycle. Immunohistochemical staining of MRL ear tissue day 7 post injury shows molecules associated with G2M. These images were overlayed onto H&E sections from the same block. Images for Wnt5a can be seen in **Fig S1CD** and for ROR2 can be seen in **Fig D1EF**. **Figures S1A, B** were reproduced from (Bedebaeva et al. 2010, PNAS).

**Figure S2.**
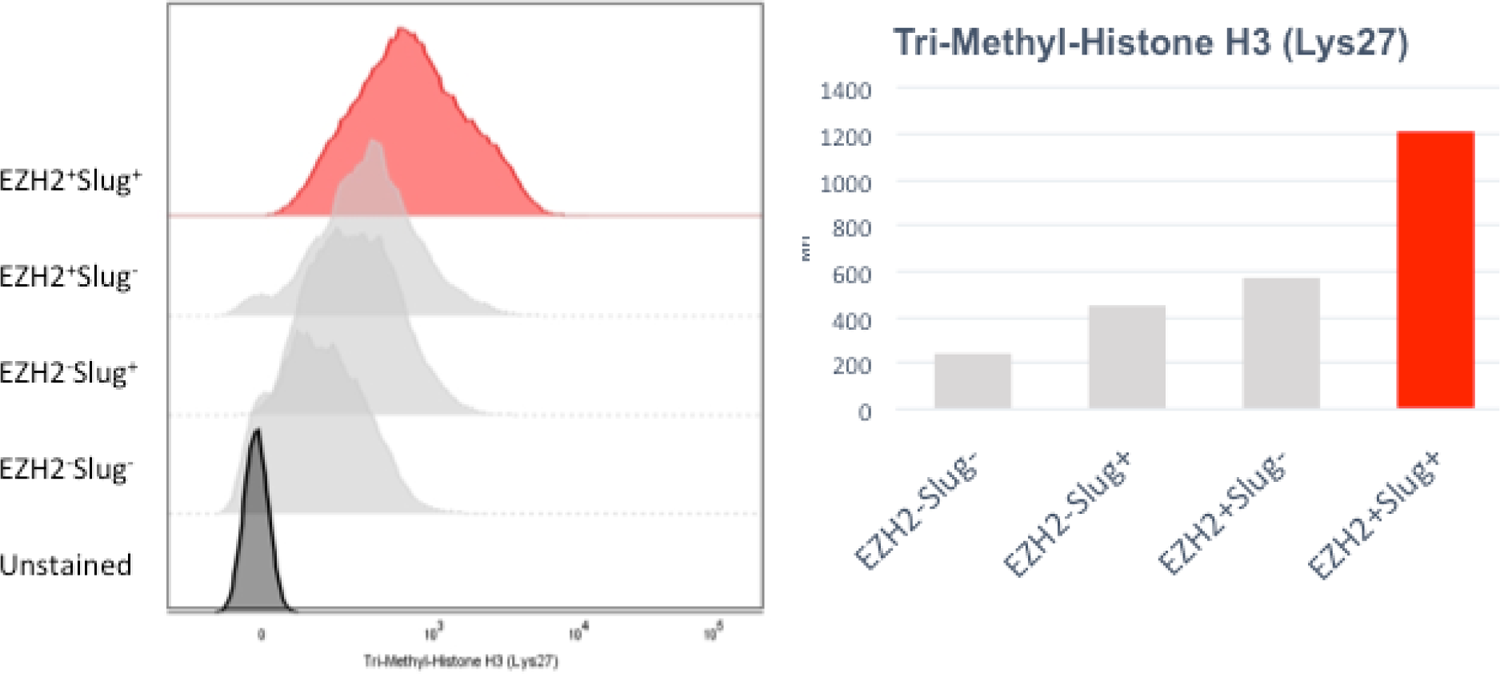
Triple staining of cells in the day 7 MRL blastema. On the left is a histogram overlay of H3K27me3 expression for all 4 populations (EZH2+Slug+, EZH+Slug-, EZH2-Slug+, EZH2-Slug-). On the right is a bar graph of the median fluorescent intensity (MFI) of H3K27me3 from each histogram on the left. The highest expression of H3K27me3 is in the EZH2+Slug+ population.

**Figure S3.**
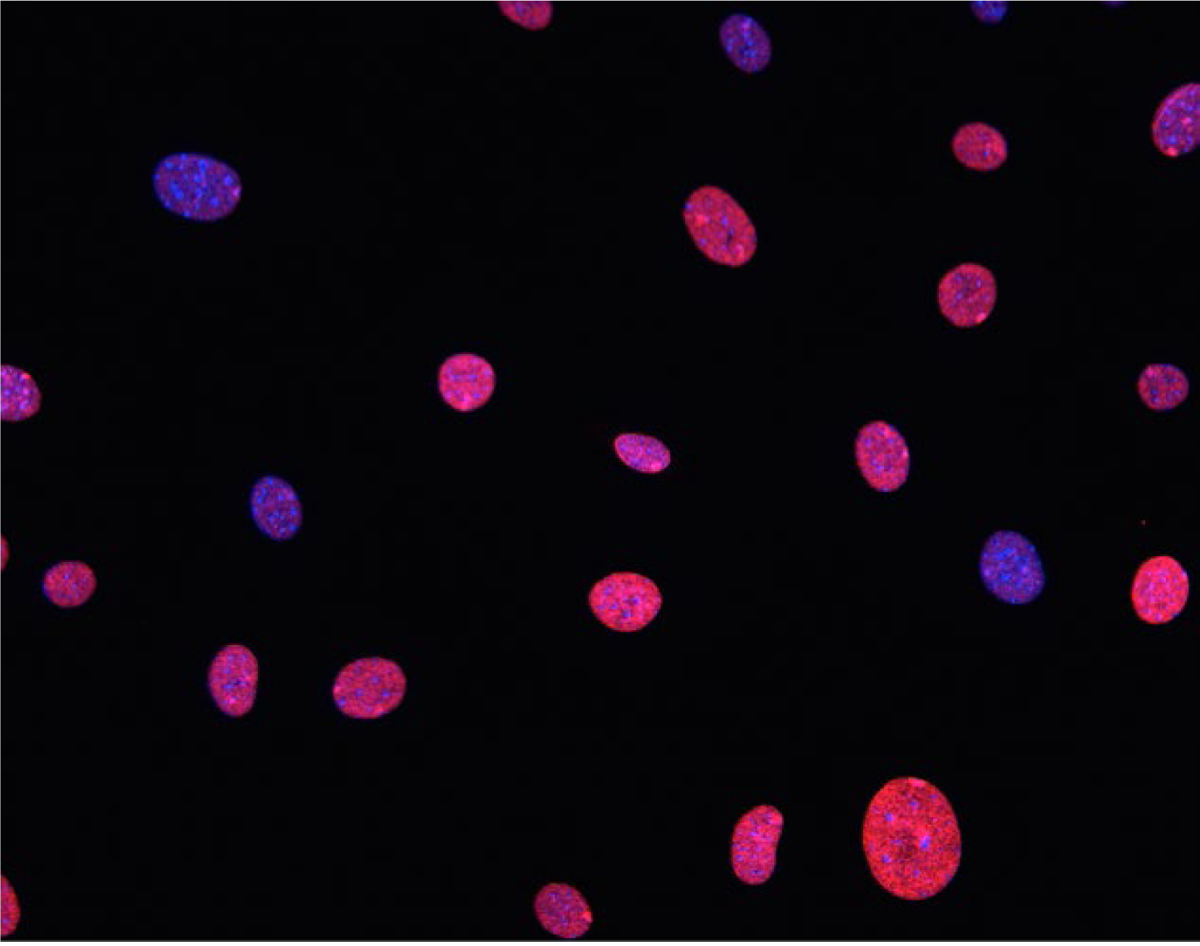
MRL-derived fibroblasts stained with antibody to H3K27me3. Seen above is a field of MRL-derived fibroblasts fixed on coverslips, stained with anti-H3K27me3, and counted. A single cell from this population is shown in Figure 8B. Over 100 cells were counted and intensely stained cells were found to be >85% positive in this population.

