## Supplementary figures and images for "Epithelial-Mesenchymal Transition: An Organizing Principle of Mammalian Regeneration"

### Supplemental Figure S1

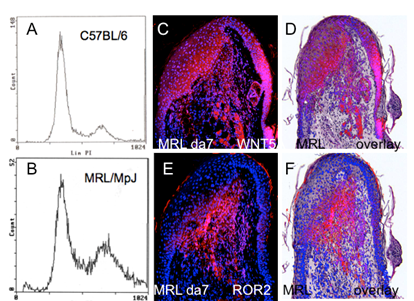

### Supplemental Figure S2

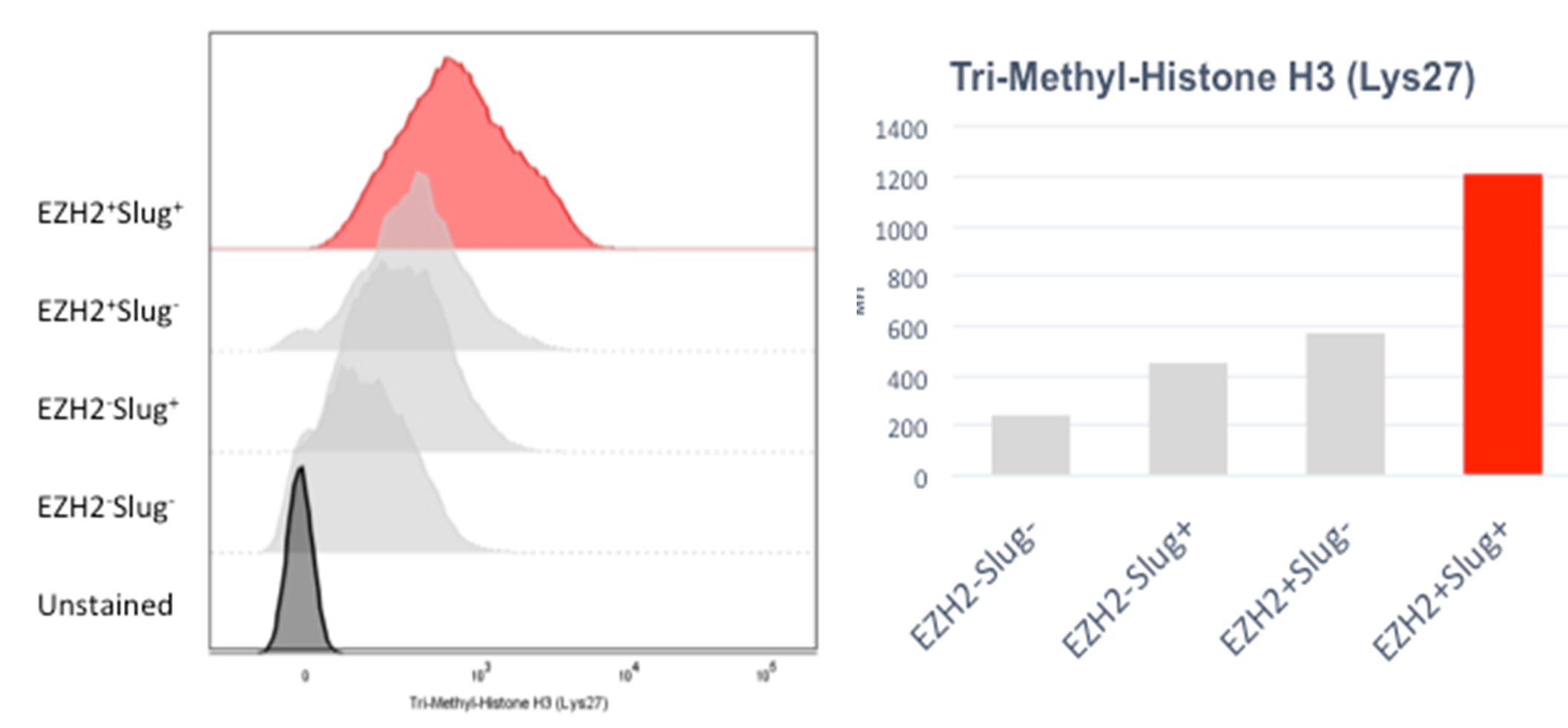

### Supplemental Figure S3

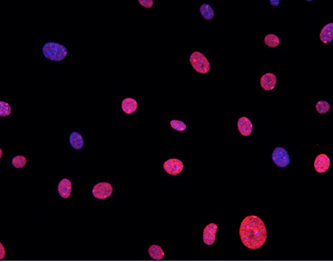
